# ZW sex chromosome structure in *Amborella trichopoda*

**DOI:** 10.1101/2024.05.10.593579

**Authors:** Sarah B. Carey, Laramie Aközbek, John T. Lovell, Jerry Jenkins, Adam L. Healey, Shengqiang Shu, Paul Grabowski, Alan Yocca, Ada Stewart, Teresa Jones, Kerrie Barry, Shanmugam Rajasekar, Jayson Talag, Charlie Scutt, Porter P. Lowry, Jérôme Munzinger, Eric B. Knox, Douglas E. Soltis, Pamela S. Soltis, Jane Grimwood, Jeremy Schmutz, James Leebens-Mack, Alex Harkess

**Author notes:** **AUTHORS FOR CORRESPONDENCE:** JLM; AH.

## Abstract

Sex chromosomes have evolved hundreds of times, and their recent origins in flowering plants can shed light on the early consequences of suppressed recombination. *Amborella trichopoda,* the sole species on a lineage that is sister to all other extant flowering plants, is dioecious with a young ZW sex determination system. Here we present a haplotype-resolved genome assembly, including highly-contiguous assemblies of the Z and W chromosomes. We identify a ∼3-Megabase sex-determination region (SDR) captured in two strata that includes a ∼300-Kilobase inversion that is enriched with repetitive sequence and contains a homolog of the *Arabidopsis* METHYLTHIOADENOSINE NUCLEOSIDASE (*MTN1-2*) genes, which are known to be involved in fertility. However, the remainder of the SDR does not show patterns typically found in non-recombining SDRs, like repeat accumulation and gene loss. These findings are consistent with the hypothesis that dioecy is recently derived in *Amborella* and the sex chromosome pair has not significantly degenerated.

## MAIN

The evolution of separate sexes, or dioecy, is a rare trait in angiosperms, having been identified in just 5-10% of species ^1^. At the same time, dioecy has evolved hundreds of independent times across the flowering plant tree of life ^2^. This makes flowering plants ideal for examining the evolution of sex chromosomes over both deep and shallow time scales. Comparative investigations of sex chromosomes rely on high-quality genome assemblies ^2^, and while the availability of genomes for dioecious species has increased, there are only a few where the structure of the sex chromosome pair has been well characterized. While divergence between X and Y sex chromosomes has been described in a growing number of angiosperm species ^2,3^, investigations of possibly less common ZW systems can shed new light on the dynamics and consequences of sex chromosome evolution.

Since its discovery as the sister lineage to all other living angiosperms, *Amborella trichopoda* (hereafter, *Amborella*) ^4–7^ has served as a pivotal taxon for investigating the origin and early diversification of flowering plants ^8,9^. *Amborella* is an understory shrub or small tree endemic to New Caledonia and the sole extant species in the Amborellales. The flowers of *Amborella* are actinomorphic and have a perianth of undifferentiated tepals, which are characteristics shared with the reconstructed ancestral flower (Fig. 1) ^9^. Importantly, however, *Amborella* is dioecious ^10^ with ZW sex chromosomes that evolved after the lineage diverged from other flowering plants ^11^. This implies that dioecy in *Amborella* is derived from a hermaphroditic mating system and that the ancestral angiosperm had perfect flowers, in agreement with ancestral state reconstructions ^9^. Significant progress has been made in several angiosperm species to identify the genes involved in the evolution of dioecy ^12–17^, but the molecular basis in *Amborella* remains unknown. Here we present a haplotype-resolved assembly of the *Amborella* genome and compare highly contiguous Z and W sex chromosome assemblies to address outstanding questions about their structure and gene content, including putative sex-determining genes.

**Fig. 1.**
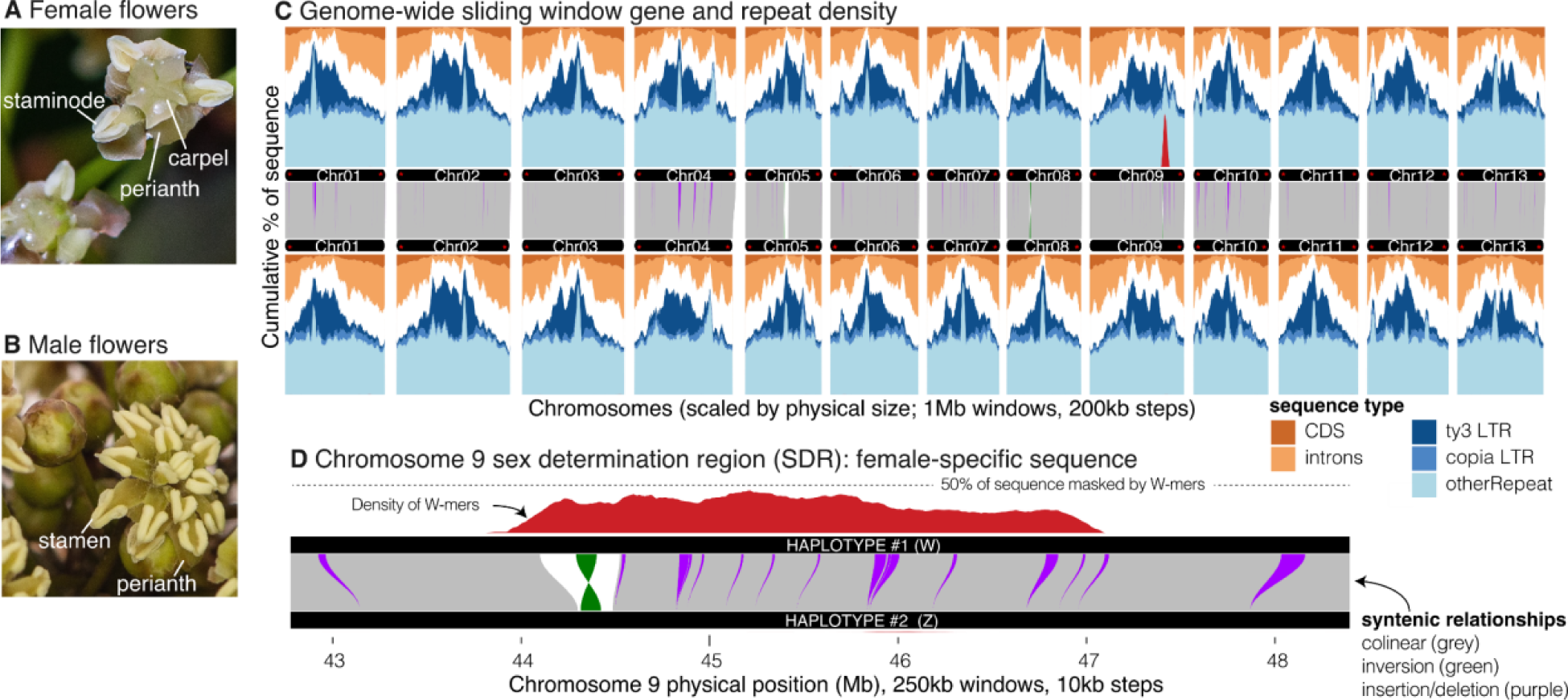
*Amborella* and its genome structure. A-B) Female and male *Amborella* flowers. The *Amborella* genome (C) and chromosome 9 (D) is typical of flowering plants: gene-rich chromosome arms and repeat-dense, large pericentromeric region. Gene positions were extracted from the protein-coding gene annotations, repeats from EDTA, and exact matches of 536,985 female-specific *k*-mers (W-mers). Syntenic mapping was calculated by AnchorWave and processed by SyRI, only plotting inversions and insertions and deletions > 10 kb. Visualization of synteny was accomplished with GENESPACE and sliding windows with gscTools. Panel B highlights the sex determination region of chromosome 9 with female-specific *k*-mers (W-mers). All chromosomes in haplotype 1 and all but four in haplotype 2 have both left and right telomeres in the assembly (flagged with red *), defined as a region of >= 150 bps made up of >= 90% plant telomere *k*-mers (CCCGAAA, CCCTAAA, RC) separated by no more than 100 bp.

## RESULTS

### Improved genome assembly and annotation of *Amborella*

The *Amborella* reference genome has been a central anchor for comparative investigations of gene family and gene structure evolution across angiosperms. Despite its demonstrated utility, the 2013 *Amborella* genome used primarily short sequencing reads, which cannot fully resolve repetitive regions ^18^. The repeat-derived gaps were filled in a more recent long-read assembly ^11^, but both biological haplotypes were collapsed into a single sequence representation. Despite the higher contiguity, the 2022 genome offers limited information regarding sex determination regions (SDRs) because in this assembly the Z and W chromosomes are a chimeric mix represented as a single chromosome ^11^.

To build a haplotype-resolved genome assembly for *Amborella* cv. Santa Cruz 75, we used a combination of PacBio HiFi (mean coverage = 58.81x per haplotype; mean read length = 22,900 bp) and Phase Genomics Hi-C (coverage = 42.31x; Table S1) sequencing technologies. The final haplotype 1 (HAP1) and 2 (HAP2) assemblies include 708.1 Mb in 59 contigs (contig N_50_ = 36.3 Mb; L_50_ = 7) and 700.5 Mb in 45 contigs (contig N_50_ = 44.5 Mb; L_50_ = 7), respectively; 99.69% and 99.87% of the assembled sequence is contained in the 13 largest scaffolds for HAP1 and HAP2, respectively, corresponding to the expected chromosome number ^19^ (Fig. S1). We found the *k*-mer completeness ^20^ of HAP1 was 95.4% (QV 63) and HAP2 was 95.3% (QV 55), and the combined assemblies exhibit 98.8% completeness (QV 57). Consistent with earlier assemblies, we annotated repeats and found they represent ∼56% of the sequence for both haplotypes (Fig. 1; Table S2) ^18^. To annotate gene models, we used a combination of RNAseq and Iso-Seq (∼757 million 2×150 read pairs, ∼825K full-length transcripts). We annotated 21,800 gene models in HAP1 and 21,721 in HAP2, with Embryophyte BUSCOs of 98.6% and 98.8%, respectively, an increase from 85.5% in the 2013 release ^18^. Overall, the new assemblies represent a great improvement in the *Amborella* genome reference, resolving most of the previous gaps (Fig. S2, Table S2).

*Amborella’*s ancient divergence ∼140 MYA ^21^ from all other living angiosperms provides an opportunity to examine conserved features that were likely present in the ancestral genome of all flowering plants. For example, the repeat-dense pericentromeric region and gene-dense chromosome arms of *Amborella* (Fig. 1) mirror those of most angiosperm genomes in stark contrast to the more uniform gene and repeat density of most conifers, ferns, and mosses ^22–24^. The pericentromeric regions are enriched in Long Terminal Repeats (LTRs), specifically *Ty3* and *Ty1* elements, as is often seen in other monocentric angiosperms ^25,26^. Interestingly, unlike many previously examined sex chromosomes, the *Amborella* Z/W do not stand out as notable exceptions in terms of gene or repeat density (Fig. 1).

### Identification of the phased *Amborella* sex chromosomes

Sex chromosomes have unique inheritance patterns relative to autosomes. In a ZW system, the non-recombining SDR of the W chromosome is only inherited by females, while the remaining pseudoautosomal region (PAR) recombines freely and is expected to show a similar lack of divergence between the sexes as the autosomes. Identification of the boundary between the SDR and PAR of sex chromosomes is nontrivial, and PAR/SDR boundaries have been shown to vary among populations in some species ^27,28^. Standard approaches for boundary identification employ combinations of methodologies like sex-biased read coverage and population genomic analyses ^29^.

To delimit the PAR/SDR boundary we performed a *k*-mer analysis ^12,30^ to identify sequences that are unique to the *Amborella* SDR (henceforth, W-mers), using four different sampling strategies (Supplemental Methods). We found the W-mers densely clustered to Chr09 at ∼44.32-47.26 Mb of HAP1 (Fig. 1-2, S3-6), supporting its identity as the W chromosome. This location is consistent with previous analyses ^11^, although we find assessing W-mers to a haplotype-resolved assembly narrows the estimated size of the SDR from ∼4 Mb to 2.94 Mb (Fig. 2, S7). Importantly, the W-mers show consistent coverage on Chr09 in HAP1, with low and sporadic coverage along any other chromosome or unincorporated scaffold in the assembly (Fig. S3-6; Table S3). In contrast to the chimeric Z/W in the previous assembly, the resulting sex chromosome assemblies are nearly complete with only four unresolved gaps in the SDR (zero gaps in the homologous region on the Z; HZR) and are fully phased (Fig. S7).

**Fig 2.**
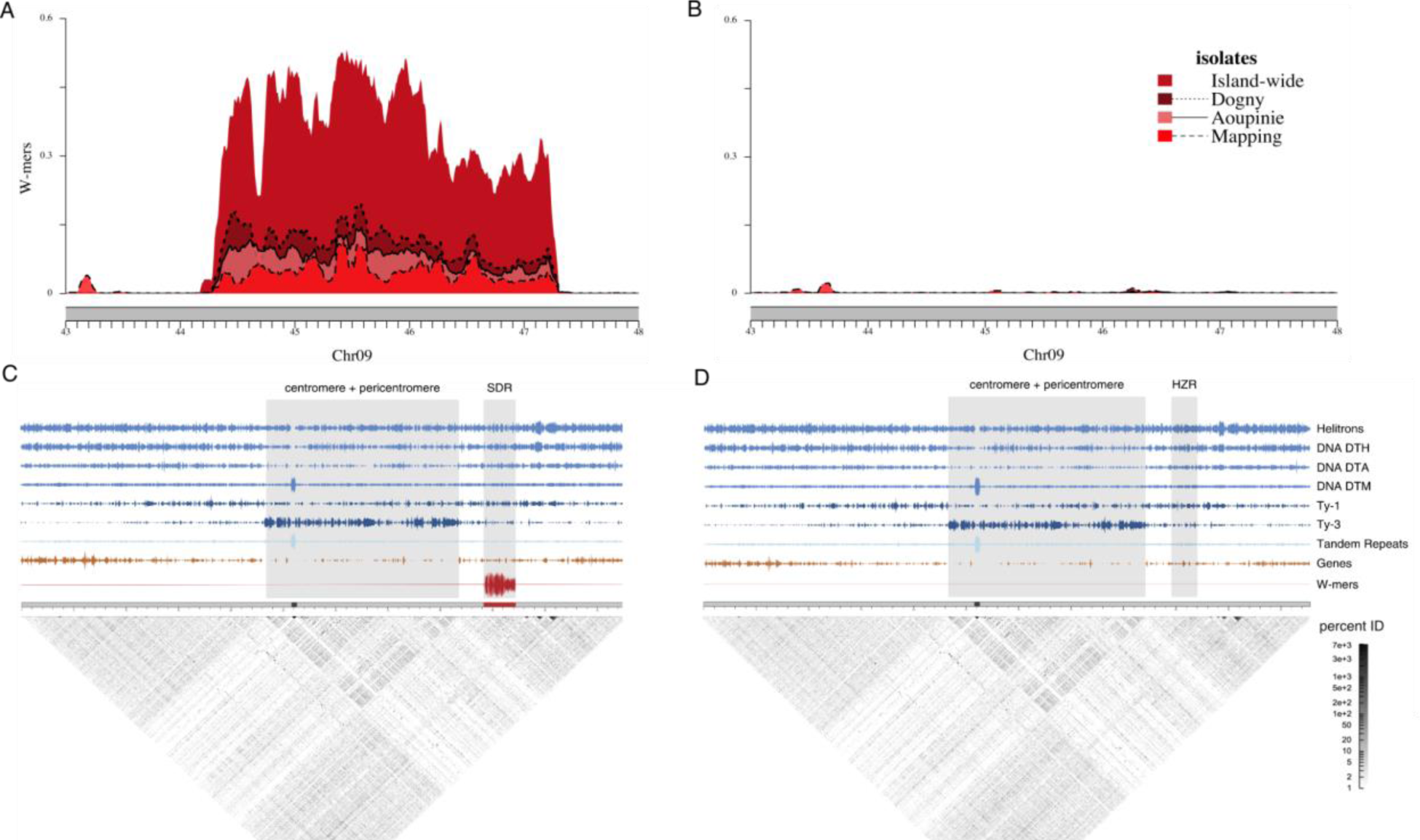
Sex chromosome location in *Amborella*. A) W-mer coverage in the sex-determining region (SDR) and (B) homologous region of the Z (HZR) using four different sampling strategies for isolates. SDR (C) and HZR (D) location and their proximity to the Chr09 centromere. Ty3 elements (dark blue) are often enriched in the pericentromeric regions of plants and correspond to the low-complexity block of tandem repeat arrays (gray) that also contain the high-complexity centromeric block, indicated by the satellite monomer density (light blue). Gene density (orange) also predictably decreases near the pericentromeric region. The SDR (red) is notably outside of the putative pericentromeric region and distant from the centromere.

A key characteristic of sex chromosomes is suppressed recombination of the SDR, and in many species, structural variants have been identified as the causal mechanism. To examine this in *Amborella*, we first used genome alignments to identify the HZR. The HZR is located on Chr09 of HAP2 at 44.52-47.12 (∼2.60 Mb; Fig. S8), suggesting the SDR is only 340 Kb larger than the HZR, which is consistent with the observed cytological homomorphy of the ZW pair ^19^. In the SDR, we found evidence for a ∼292-Kb inversion located ∼20 Kb within the beginning of the boundary and containing the majority of the W-specific sequence (Figs. 1B, S9). We could not, however, find evidence for inversions or other large structural variants surrounding the remaining portion of the SDR. Instead, the Z and W chromosomes are highly syntenic with one another, similar to the autosomes (Figs. 1, S9). We investigated other potential mechanisms for suppressed recombination, such as proximity to centromeres, where the existing low recombination has been shown to facilitate SDR evolution in some species ^31^. In *Amborella*, the SDR is not located near the centromere; rather, it is approximately 1.82 Mb away from the *Ty3*-retrotransposon-rich pericentromeric region (Fig. 2). In the absence of obvious structural variants encompassing the SDR, it suggests that *Amborella* has a non-canonical mechanism to enforce non-recombination between the Z and the W.

### The *Amborella* sex chromosomes are evolutionarily young

*Amborella*’s sex chromosomes have previously been shown to have evolved after the lineage split from other living flowering plants ^11^. With our phased Z/W pairs, we can better determine Z- and W-linked genes, providing a more confident estimate of the age of the SDR, and examine gene gain events. A classic signature of multiple recombination suppression events is a stepwise pattern of synonymous substitutions (Ks), where genes captured into the SDR in the same event are expected to have similar levels of Ks (i.e., strata) and the oldest captures have the highest Ks values ^32^. Understanding this timing of gene gain is essential to understanding the genetic mechanism for sex determination, because the candidate sex-determining genes are likely to have ceased recombining first (barring turnovers ^29^).

To examine gene gain in the *Amborella* SDR, we calculated Ks of one-to-one orthologs on the W and Z chromosomes (i.e., gametologs). We compared the Ks values of 45 identifiable gametologs to 1,397 one-to-one orthologs in the PARs. We found that Ks varies across the SDR-HZR portion of the sex chromosomes (0.002-0.20; mean Ks=0.0298, SD=0.032) and is significantly higher than Ks in the PARs (mean Ks=0.004, SD=0.019; Kruskal-Wallis p<0.00001); Fig. S10), consistent with the expectation that the SDR is diverging from the HZR on the Z chromosome. Interestingly, the gametolog pair with the highest Ks within the SDR is a homolog of *Arabidopsis* METHYLTHIOADENOSINE NUCLEOSIDASE *MTN1-2*, a gene involved in fertility, suggesting it resides in the oldest portion of the SDR; notably, the location of the W-linked *MTN1-2* homolog is within the SDR inversion.

We found evidence for two strata of gene capture into the SDR (Fig. 3). The Ks values show two distinct steps, with the higher Ks values in the region corresponding to the inversion. Defining the precise boundary of strata without obvious structural variants can be a challenge. To delineate stratum one (S1) from two (S2), we assessed W-mer density and the average nucleotide differences between sampled females and males (Nei’s dXY). We found the drop in W-mers and dXY in sliding windows coincides with a drop in dXY when run on only the gene models (Fig. 3). Using this line as our boundary between strata, we found dXY of genes to be significantly different (Mann Whitney U, p<3e-7), higher in S1 (mean = 0.0167, n=62) than S2 (mean = 0.0081, n=35; entire Chr09 = 0.0038; n=1908). We also found Ks to be significantly different between the strata (S1 mean Ks=0.037, SD=0.037; S2 mean Ks=0.021, SD=0.023; Mann-Whitney U, p=0.0014) as was the extent of nonsynonymous changes in proteins (Ka; Mann-Whitney U, p=0.008; Fig. 3), supporting inference of two strata. Using Ks, we also estimated the age of the SDR in *Amborella*. Following the previously applied approach ^11^, we found S1 to be ∼4.97 MYA while S2 is nearly half as old at ∼2.41 MYA. These analyses indicate that the *Amborella* sex chromosomes are evolutionarily young, similar to several well-characterized XY systems ^3^, and further suggest that the sex chromosomes evolved well after the lineage split from the rest of angiosperms.

**Fig. 3.**
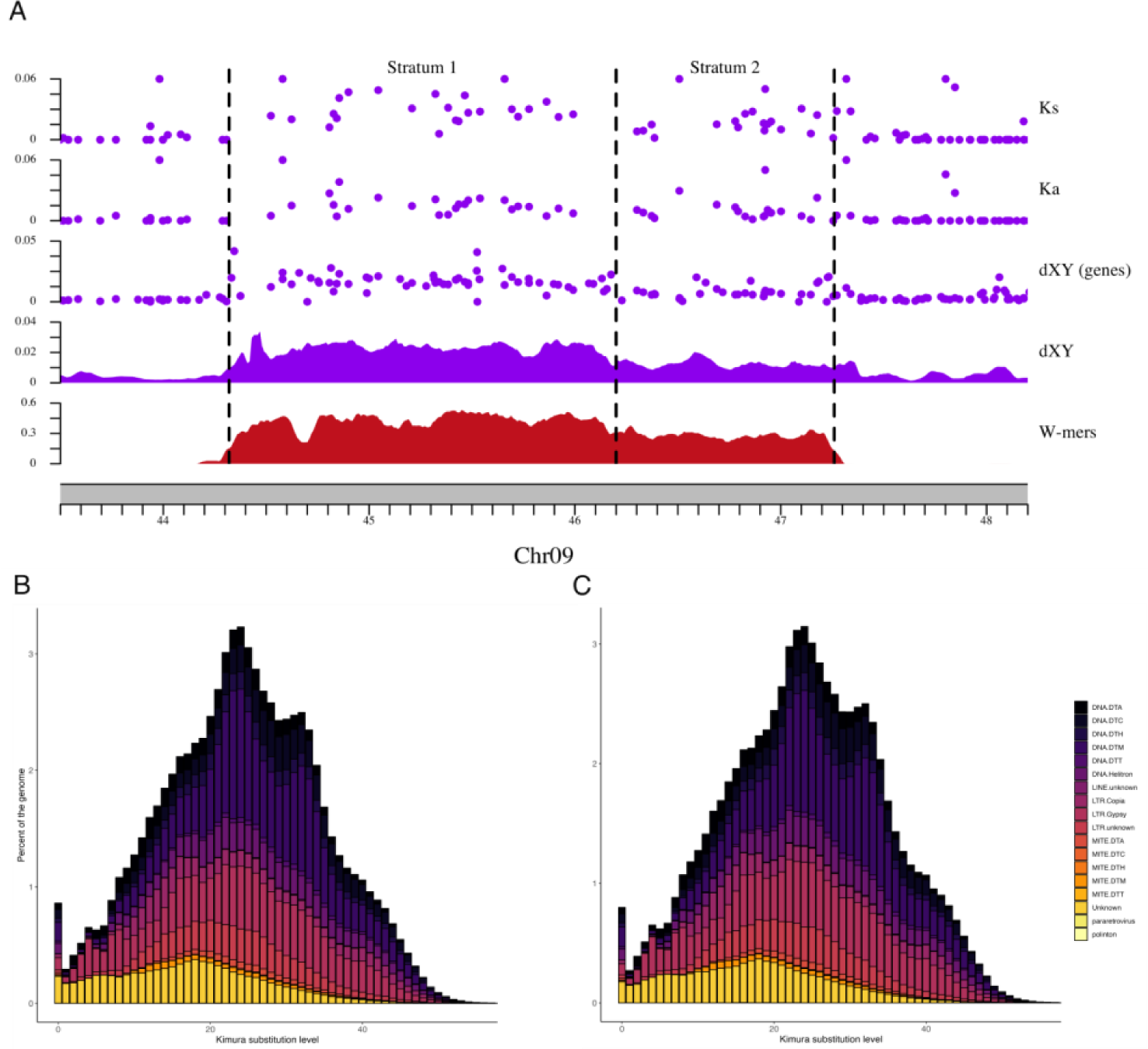
Molecular evolution of the *Amborella* sex chromosomes. A) Evidence for two strata. Points above 0.06 were excluded. B-C) The repeat landscapes of the *Amborella* haplotypes indicate similar patterns of expansion and minimal evidence of recent TE proliferation. Relative time is determined by the Kimura substitution level with lower values closer to 0 representing more recent events and higher values approaching 40 representing older events.

### The *Amborella* W shows little degeneration

The recent origin of the *Amborella* sex chromosomes provides an opportunity to examine the early stages of their evolution. The lack of recombination in an SDR reduces the efficacy of natural selection and drives the accumulation of slightly deleterious mutations ^33,34^. Two parallel signatures of deleterious mutations seen across independent evolutions of sex chromosomes is the accumulation of repeats and the loss of genes ^35–38^. However, the tempo of this process of degeneration is not well understood.

In the SDR of *Amborella,* curiously we overall do not find the expected patterns of repeat expansions found in other SDRs. At 51.66% repeat elements, the SDR is lower than the genome average (56%) and 0.05% lower than the HZR. The only observed enrichment in repeats is within the inversion, where we find more *Ty3* LTRs (4.32% increase relative to the HZR; Fig. 2). Otherwise, only a slight distinction between the SDR and its HZR is evident: the SDR exhibits a marginal increase ranging between 0.01-0.13% in the density of some superfamily elements (Fig. 2; Table S4). We examined the distribution of the divergence values for intact LTRs as a proxy for their age ^39^ but found no patterns of distinctly younger or older LTRs within the W or Z (Fig. S11). Moreover, to assess genome-wide repeat expansion across the major Transposable Element (TE) superfamilies ^40^, we used repeat landscapes, which showed a comparable pattern within the Z/W (Fig. 3, S12). These observations support previous characterization of TE insertions in the *Amborella* genome as being quite old with little proliferation over the last 5 MYA^18^. It has been proposed that a loss of active transposases or silencing may be playing a role in reducing TE activity across the *Amborella* genome ^18^ including the SDR.

Gene loss in an SDR has been hypothesized to contribute to the evolution of heteromorphy seen in many sex chromosome pairs ^41,42^. In *Amborella*, of the 97 annotated models in the SDR and 84 in the HZR, 37 were W-specific and 24 Z-specific. To examine whether these models were missing from the other haplotype for technical or biological reasons, we also used dXY and presence-absence variation (PAV) between the sexes to evaluate gene content. For most of the W-specific models, males showed presence, and dXY within females was comparable to that of identifiable gametologs (mean dXY = 0.0136; Table S5). Only seven models showed absence in coverage in males (dXY = 0 in females), suggesting conservatively that these represent W-specific genes, four of which are in the SDR inversion. Similarly, we identified only six Z-specific gene models. These analyses suggest that the Z and W have similar numbers of haplotype-specific genes and that the SDR has experienced similar levels of gene loss as the HZR.

Together, these results provide little evidence that degenerative processes, associated with cessation of recombination, have occurred in the *Amborella* SDR. This region is younger than that of *Rumex* (5-10 MYA ^43^) and *Silene* (10 MYA ^44^), which both show signatures of degeneration ^38,45^. However, in *Spinacia oleracea*, a younger SDR (2-3 MYA) does show signs of degeneration ^46,47^. The tempo of degeneration is apparently slower in *Amborella* and there has not been sufficient time for gene loss or an accumulation of repeats as a consequence of the loss of recombination.

### Candidate sex-determining genes in *Amborella*

ZW sex chromosomes have been less well-characterized in plants than in animals; thus, *Amborella* can provide unique insights regarding the genetic mechanisms associated with their evolution. The two-gene model for sex chromosome evolution associated with a transition from hermaphroditism to dioecy posits that distinct genes with antagonistic impacts on female and male function experience strong selection for tight linkage (i.e., loss of recombination)^48^. Under this model, evolution of a ZW sex chromosome pair requires a dominant mutation causing male sterility arising on a proto-W chromosome, followed by a recessive loss-of-female-function mutation on the proto-Z (assuming a gynodioecious intermediate) ^48^. Identification of these sex-determining genes relies on an understanding of when sterility arises in the carpel and stamen developmental pathways. In *Amborella,* ontogenetic differences between female and male flowers are seen early in development. Whereas male flowers produce an average of 12 stamens spiraling into the center of the flower, female flowers typically initiate a few staminodes just inside the tepals, but carpel initiation replaces stamenoid initiation as organ development proceeds towards the center of the flower ^49^ (Fig. 1).

To identify candidate sex-determining genes, we examined differential expression between female and male flower buds during stage 5/6 of flower development, when carpels, stamens, and microsporangia develop ^11,49,50^. We found 1,777 significantly differentially expressed genes at an adjusted p-value greater than 0.05. Of these, 34 are in the SDR, several of which are well-known flower development genes, including homologs of *MTN1-2*, *WUSCHEL (WUS)*, *LONELY GUY (LOG)*, *MONOPTEROS/Auxin Response Factor 5 (MP/ARF5)*, and *small auxin up-regulated RNA* (*SAUR*) gene families (Fig. S13; Table S6-7). We found *ambMTN* and *ambLOG* had higher transcript abundance in females, while *ambWUS*, *ambMP*, and *ambSAUR* had greater expression in males. To further examine the sex-specific expression of SDR genes, we used the EvoRepro database (https://evorepro.sbs.ntu.edu.sg/), which has transcriptome data for 16 different tissue types for *Amborella* ^51^. We contrasted female and male buds and flowers and found three genes with male-biased transcript abundance: *ambWUS* and a *DUF827* gene in buds and *ambLOG* in flowers, the latter differing in which sex has higher abundance from the analyses using stage 5/6 flowers. Given the known functions of these genes in *Arabidopsis* flower development, they are strong candidates for investigation of sex determination in *Amborella*.

While functional analyses are not currently possible in *Amborella*, comparisons to other species implicate the function of candidate genes that may be playing roles in *Amborella* sex determination. *WUS* is a homeobox transcription factor that is required for the maintenance of the floral meristem and has been shown to influence gynoecium and anther development ^52,53^ In *Arabidopsis*, knockouts have sepals, petals, a single stamen, and no carpel ^54^. *WUS* has also been implicated in sex determination or shown sex-specific expression in several species that have unisexual flowers. In monoecious castor bean (*Ricinus*), *WUS* expression was only found in the shoot apical meristem of male flowers ^55^, and in cucumbers (*Cucumis*), *WUS* expression is three times greater in the carpel primordia of male flowers than females ^56^. In *Silene*, gynoecium suppression is controlled by the *WUSCHEL-CLAVATA* feedback loop ^16^. Interestingly, we do not see male-biased expression of the *CLV3* ortholog in *Amborella,* but we do see female-biased transcript abundance of the *Amborella CLE40* ortholog. In *Arabidopsis*, *WUS* promotes *CLV3* expression in the central zone of the inflorescence meristem while suppressing *CLE40* expression in the peripheral zone ^57^. It is possible that the smaller floral meristem seen in female development relative to male floral meristems is due to reduced *ambWUS* expression driving increased *ambCLE40* expression and encroachment of peripheral zone cells into the central zone of the floral meristem. The role of *WUS* in maintaining meristematic zonation, coupled with its position in S1 in the SDR, makes *ambWUS* a strong candidate for playing some role in gynoecium suppression. Another strong candidate is *ambLOG*. *LOG* mutants were originally characterized in rice as producing floral phenotypes with a single stamen and no carpels ^58^; in date palms (*Phoenix*), a *LOG*-like gene was identified as a candidate Y-chromosome-linked female suppression gene ^13^. In *Amborella*, *ambLOG* showed greater expression in females in the stage 5/6 data but was male-biased based when considering all 16 tissues in the EvoRepro dataset. This switch in sex bias, and the fact *ambLOG* is located in the younger stratum of SDR (S2), suggest that differential *ambWUS* (and *ambCLE40*) expression may have been a first step in the divergence of male and female flower development. Like *ambLOG*, the *ambMP* and *ambSAUR* genes were captured in S2, and their functions in *Arabidopsis* suggest other roles in sex-specific development. *MP* has been shown to be involved with apical patterning of the embryo axis ^59,60^. *SAUR*s are a large gene family and in general play a role in cell elongation ^61^, including in pollen tube growth ^62^, stamen filament elongation ^63^, and pistil growth ^64^. Without functional validation in *Amborella*, we cannot rule out the possibility of any of these genes, though based on the data available, *ambWUS* may be the strongest candidate for spurring divergence in male and female flower development.

The significant difference in gene expression of *ambMTN* is especially interesting given that it is the gene model with the highest Ks value that is located in the SDR inversion. *MTN1-2* genes encode 5’-methylthioadenosine (MTA) nucleosidase ^65^, and double mutant *mtn1-1mtn2-1* flowers in *Arabidopsis* have indehiscent anthers and malformed pollen grains ^66^. Double mutants also affected carpels and ovules, although the structures were aberrant but not necessarily non-functional, and 10% looked like wild type ^66^. The observed anther phenotype in *Arabidopsis* is consistent with the staminode development in female flowers in *Amborella*, and together these lines of evidence suggest that *ambMTN* may be the male-sterility gene. Based on our analyses, we hypothesize that the W-linked *ambMTN* was the initial male-sterility mutation, creating the proto-W, followed by a loss-of-function mutation on the W-*ambWUS* and a Z-copy shift to dosage dependant gynoecium suppression. In sum, we hypothesize that *Amborella* follows the two-gene model for sex chromosome evolution and dioecy. The genes we have identified here make ideal candidates for further functional genomic investigation and validation.

## DISCUSSION

Recent advances in sequencing technologies and assembly algorithms have enabled the construction of telomere-to-telomere genome assemblies for humans, including the X and Y sex chromosomes ^67,68^. The sex chromosomes in humans and other animals are often highly heteromorphic and can be the most challenging chromosomes to sequence and assemble ^69^. Moreover, given their antiquity, it is not possible to reconstruct events dating back to the origin and early evolution of mammalian sex chromosomes. Plants, however, have repeatedly evolved sex chromosomes derived from different ancestral autosomes, with different sex-determining mutations ^2,3^ and with various mechanisms to impede recombination between the sex chromosome pair. Here we show that we can fully phase structurally similar sex chromosomes within a heterogametic individual. Our analyses highlight the utility of phased sex chromosomes, and diversity sequencing, to develop models of sex chromosome evolution when experimental investigation of gene function is currently intractable. This research lays the foundation for examining sex chromosome evolution in all angiosperms, starting with the sister species to all living flowering plants, *Amborella*.

## Acknowledgements

The work (proposal no. 10.46936/10.25585/60001405) conducted by the U.S. Department of Energy (DOE) Joint Genome Institute (https://ror.org/04xm1d337), a DOE Office of Science User Facility, is supported under contract no. DE-AC02-05CH11231. Additional support for analysis was provided by the United States Department of Agriculture National Institute of Food and Agriculture Postdoctoral Fellowship no. 2022-67012-38987 (S.B.C.), National Science Foundation (NSF) IOS-PGRP CAREER no. 2239530 (A.H.), and National Science Foundation GRFP (L.A.). We thank the Atlanta Botanical Garden for providing *Amborella* material used in this study and Adam Bewick for the images of *Amborella* flowers.

## Author contributions

Concept and research design: S.B.C., J.S., J.L.-M., A.H.

Sample collection, data collection, sequencing: A.S., T.J., K.B., P.P.L., J.M., E.B.K., D.E.S., P.S.S., J.G., J.L.-M.

Genome assembly and annotation: S.B.C., J.J., S.S.

Computational and statistical analyses: S.B.C., L.A., J.T.L., A.L.H., P.G., A.Y.

Wrote the paper (with contributions from all authors): S.B.C., L.A., J.T.L., J.J., A.L.H., C.S., D.E.S., P.S.S., J.L.-M., A.H.

## Data availability

The genome assemblies and annotations (v.2.1) are available on Phytozome (https://phytozome-next.jgi.doe.gov/) and have been deposited on NCBI under XXXX. Sequencing libraries are publicly available on NCBI under BioProject PRJNA1100625. Individual accession numbers are provided in Supplementary Table S8-9.

## MATERIALS AND METHODS

### DNA/RNA extraction, library prep, and sequencing

We sequenced *Amborella trichopoda* (var. Santa Cruz 75) using a whole genome shotgun sequencing strategy and standard sequencing protocols. High molecular weight DNA was extracted from young tissue using the protocol of Doyle and Doyle ^70^ with minor modifications. Flash-frozen young leaves were ground to a fine powder in a frozen mortar with liquid nitrogen followed by very gentle extraction in a 2% CTAB buffer (that included proteinase K, PVP-40 and beta-mercaptoethanol) for 30 minutes to 1 hour at 50 °C. After centrifugation, the supernatant was gently extracted twice with 24:1 Chloroform : Isoamyl alcohol. The upper phase transferred to a new tube and added 1/10th volume of 3 M Sodium acetate, gently mixed, and DNA precipitated with iso-propanol. DNA precipitate was collected by centrifugation, washed with 70% ethanol, air dried for 5-10 minutes and dissolved thoroughly in an elution buffer at room temperature followed by RNAse treatment. DNA purity was measured with a Nanodrop, DNA concentration measured with Qubit HS kit (Invitrogen, Waltham, MA) and DNA size was validated by CHEF-DR II system (Bio-Rad Laboratories, Hercules, CA). The PacBio HiFi libraries were sequenced at the HudsonAlpha Institute for Biotechnology in Huntsville, Alabama. The PacBio HiFi library was constructed using DNA that was sheared using a Diagenode Megaruptor 3 instrument. Libraries were constructed using SMRTbell Template Prep Kit 2.0 and tightly sized on a SAGE ELF instrument (1-18kb) to a final library average insert size of 24k. Sequencing was completed using the SEQUEL II platform. For the PacBio sequencing, a total raw sequence yield of 83.3 Gb, with a total coverage of 58.81x per haplotype (Table S10).

The Illumina Hi-C reads for Santa Cruz 75 were sequenced at Phase Genomics with a single 2×80 Dovetail Hi-C library (169.27x; Table S1). The Illumina PCR-free library was extracted using a Qiagen DNeasy kit (Qiagen, Hilden, Germany) and was sequenced at the HudsonAlpha Institute for Biotechnology in Huntsville, Alabama. Illumina reads were sequenced using the Illumina NovaSeq 6000 platform using a 400 bp insert TruSeq PCRfree fragment library (49.62×). Prior to assembly, Illumina fragment reads were screened for phix contamination. Reads composed of >95% simple sequence and those <50 bp after trimming for adapter and quality (q<20) were removed. The final read set consists of 158,007,088 reads for a total of 49.62× of high-quality Illumina bases.

To annotate gene models, we generated RNAseq and Iso-Seq data on several stages of leaf, flower, and fruit for Santa Cruz 75 and two male isolates, ABG 2006-2975 and ABG 2008-1967 (Table S8). Total RNA were extracted using a Qiagen RNeasy kit. The PacBio Iso-Seq libraries were constructed using a PacBio Iso-Seq Express 2.0 kit. Libraries were either sized (0.66x bead ratio) or unsized (1.2× bead ratio) to give final libraries with average transcript sizes of 2kb or 3kb respectively. Libraries were sequenced using polymerase V2.1 on a PacBio Sequel II Platform. The RNASeq libraries were constructed using an Illumina TruSeq Stranded mRNA Library Prep Kit using standard protocols. Libraries were sequenced using a NovaSeq 6000 Instrument PE150 to 40 million reads per library.

To identify the sex chromosomes, we additionally whole-genome sequenced 52 *Amborella* isolates (Table S9). DNA extractions were performed using a standard CTAB protocol. Illumina sequencing was performed on NovaSeq and HiSeq platforms at RAPiD Genomics in Gainesville, Florida using a 2×150 paired end library. The voucher specimens are deposited at the New Caledonia Herbarium in Nouméa (Herbarium code: NOU) and Indiana University (IND). Existing data used to support this manuscript are found in Table S9.

### Genome assembly

The version 2.0 HAP1 and HAP2 assemblies were generated by assembling the 3,605,703 PacBio CCS reads (58.81x per haplotype) using the HiFiAsm+HIC assembler ^71^ and subsequently polished using RACON ^72^. This produced initial assemblies of both haplotypes. The HAP1 assembly consisted of 1,522 scaffolds (1,522 contigs), with a contig N50 of 25.5 Mb, and a total genome size of 800.6 Mb (Table S11). The HAP2 assembly consisted of 1,043 scaffolds (1,043 contigs), with a contig N50 of 43.0 Mb, and a total genome size of 773.5 Mb (Table S11).

Hi-C Illumina reads from *Amborella trichopoda* (var. Santa Cruz 75), were separately aligned to the HAP1 and HAP2 contig sets with Juicer ^73^, and chromosome scale scaffolding was performed with 3D-DNA ^74^. No misjoins were identified in either the HAP1 or HAP2 assemblies. The contigs were then oriented, ordered, and joined together into 13 chromosomes per haplotype using the HiC data. A total of 31 joins was applied to the HAP1 assembly, and 20 joins for the HAP2 assembly. Each chromosome join is padded with 10,000 Ns. Contigs terminating in significant telomeric sequence were identified using the (TTTAGGG)_n_ repeat, and care was taken to make sure that they were properly oriented in the production assembly. The remaining scaffolds were screened against bacterial proteins, organelle sequences, GenBank nr and removed if found to be a contaminant. After forming the chromosomes, it was observed that some small (<20Kb) redundant sequences were present on adjacent contig ends within chromosomes. To resolve this issue, adjacent contig ends were aligned to one another using BLAT ^75^, and duplicate sequences were collapsed to close the gap between them. A total of 5 adjacent contig pairs were collapsed in the HAP1 assembly and 4 in the HAP2 assembly.

Finally, homozygous SNPs and INDELs were corrected in the HAP1 and HAP2 releases using ∼49x of Illumina reads (2×150, 400bp insert) by aligning the reads using BWA-MEM ^76^ and identifying homozygous SNPs and INDELs with the GATK’s UnifiedGenotyper tool ^77^. A total of 465 homozygous SNPs and 15,763 homozygous INDELs were corrected in the HAP1 release, while a total of 473 homozygous SNPs and 17,208 homozygous INDELs were corrected in the HAP2 release. The final version 2.0 HAP1 release contained 707.9 Mb of sequence, consisting of 59 contigs with a contig N50 of 36.3 Mb and a total of 99.69% of assembled bases in chromosomes. The final version 2.0 HAP2 release contained 700.3 Mb of sequence, consisting of 45 contigs with a contig N50 of 44.5 Mb and a total of 99.87% of assembled bases in chromosomes.

### Genome annotation

Transcript assemblies were made from ∼757M pairs of 2×150 stranded paired-end Illumina RNAseq reads using PERTRAN, which conducts genome-guided transcriptome short read assembly via GSNAP ^78^ and builds splice alignment graphs after alignment validation, realignment and correction. To obtain 825K putative full-length transcripts, about 20M PacBio Iso-Seq CCSs were corrected and collapsed by a genome-guided correction pipeline, which aligns CCS reads to the genome with GMAP ^78^ with intron correction for small indels in splice junctions, if any, and clusters alignments when all introns are the same or 95% overlap for single exon. Subsequently 563,694 transcript assemblies were constructed using PASA ^79^ from ESTs and RNAseq transcript assemblies described above. Loci were determined by transcript assembly alignments and/or EXONERATE alignments of proteins from *Arabidopsis thaliana*, *Glycine max*, *Sorghum bicolor*, *Oryza sativa*, *Lactuca sativa*, *Helianthus annuus*, *Cynara cardunculus*, *Selaginella moellendorffii*, *Physcomitrella patens*, *Nymphaea colorata*, *Solanum lycopersicum*, and *Vitis vinifera,* and Swiss-Prot eukaryote proteomes to the repeat-soft-masked *Amborella trichopoda* HAP1 genome using RepeatMasker ^80^ with up to 2K BP extension on both ends unless extending into another locus on the same strand. Gene models were predicted by homology-based predictors, FGENESH+ ^81^, FGENESH_EST (similar to FGENESH+, but using EST to compute splice site and intron input instead of protein/translated ORF), EXONERATE ^82^, PASA assembly ORFs (in-house homology-constrained ORF finder), and AUGUSTUS ^83^ trained by the high-confidence PASA assembly ORFs and with intron hints from short-read alignments. The best scored predictions for each locus were selected using multiple positive factors, including EST and protein support, and one negative factor: overlap with repeats. The selected gene predictions were improved by PASA and the optimal set was selected using several curated gene quality metrics ^84^. We assessed the gene annotations using compleasm v0.2.6 ^85^ using the Embryophyta database.

We further annotated repeats using EDTA v2.0.0 ^86^ using the sensitive mode that runs RepeatModeler ^87^. To identify tandem repeats, we used Tandem Repeats Finder ^88^ (parameters 2 7 7 80 10 50 500 -f -d -m -h). We ran StainedGlass v0.5^89^ to visualize the massive tandem repeat arrays for chromosomes in both haplotypes. To build the repeat landscapes for assessing recent expansion events, we followed the methods outlined in EDTA Github Issue #92: Draw Repeat Landscapes, utilizing a library generated from an independent annotation on the combined haplotypes with EDTA v2.0.1.

### Comparisons between assembly haplotypes

To plot comparisons between the two haplotypes, including genes and repeats, we used GENESPACE v.1.3.1 ^90^. To generate synteny between the two haplotypes, we first performed genome alignments. Haplotype 1 and haplotype 2 were aligned using AnchorWave ^91^ using the ‘genoAli’ method and ‘-IV’ parameter to allow for inversions. Alignment was performed using only “chromosome” sequence for each haplotype. The alignment was converted to SAM format using the ‘maf-convert’ tool provided in ‘last’ ^92^ and used for calling variants with SyRI ^93^. The output from SyRI was used to make chromosome-level synteny and SV plots using plotsr ^94^.

### Identification of the sex chromosome non-recombining region

We used whole-genome sequencing data to identify the sex-determining region (SDR) of the W. All paired-end Illumina data had adapters removed and were quality filtered using TRIMMOMATIC v0.39 ^95^ with leading and trailing values of 3, sliding window of 30, jump of 10, and a minimum remaining read length of 40. We next found all canonical 21-mers in each isolate using Jellyfish v2.3.0 ^96^ and used the bash *comm* command to find all *k*-mers shared in all female isolates and not found in any male isolate (W-mers). We mapped the W-mers to both haplotype assemblies using BWA-MEM v0.7.17 ^76^, with parameters ‘-k 21’ ‘-T 21’ ‘-a’ ‘-c 10’. W-mer mapping was visualized by first calculating coverage in 100,000-bp sliding windows (10,000 bp jump) using BEDTools v2.28.0 ^97^ and plotted using karyoploteR v1.26.0 ^98^.

### Structural variation

To identify structural variants between the haplotypes, we mapped PacBio reads using minimap2 v2.24 ^99^ in HiFi mode, added the MD tag using samtools v1.10 *calmd*, and called structural variants using Sniffles v2.0.7 ^100^. We also performed whole genome alignments using minimap2 v2.24 ^99^ and visualized the dotplot using pafR v0.0.2 ^101^.

### Gene homology and protein evolution

To identify one-to-one orthologs on the ZW to examine protein evolution, we ran OrthoFinder v.2.5.2 ^102,103^ using only the *Amborella* haplotypes. We calculated synonymous (Ks) and nonsynonymous (Ka) changes in codons using Ka/Ks Calculator v2.0 ^104^.

### Nucleotide differences between the sexes

BWA v0.7.17 ^76^ was used to map reads and bcftools v1.9 *mpileup* and *call* ^105^ functions were used to call variants using the Island-wide sampling (nine male and six female plants; Table S9). We filtered the vcf file using ‘QUAL>20 & DP>5 & MQ>30’, minor allele frequency of 0.05, and dropped sites with > 25% missing data. To calculate Nei’s nucleotide diversity between the sexes (dXY) we used pixy v1.2.7.beta1 ^106^. dXY was calculated using 100,00bp windows with a 10,000bp jump, and on the gene models only separately.

### Presence-absence variation

Presence-absence variation (PAV) was identified following the methods of Hu et al. ^107^ mapping reads from the Island-wide sampling (eight male and six female plants; the Atlanta Botanical Gardens isolate was removed due to low resequencing depth; Table S9) to our new reference genome and annotation. Briefly, reads for the samples were aligned to each haplotype using BWA v0.7.17 ^76^. Sorted BAM files were converted to bedgraph format using bedtools v2.30.0 ^97^. Genes were called absent if the horizontal coverage of exons was <5% and the average depth was <2×. A test for equality in the proportion of PAV rate across chromosomes was performed in R using the ‘prop.test()’ function.

### Gene expression analyses

To examine gene expression and identify candidate sex-determining genes, we used existing RNAseq data from 10 females and 10 males ^11^. From the reads, we first filtered using TRIMMOMATIC (same parameters as above). Filtered reads were mapped to the haplotype 1 genome assembly using STAR v2.7.9a ^108^ and expression estimated for the annotated gene models using StringTie v2.1.7 (-e, -G) ^109^. We performed differential gene expression analyses using DESeq2 v1.32.0 ^110^, with the contrast being between the sexes.

