## Supplemental Materials for "ZW sex chromosome structure in *Amborella trichopoda*"

### SUPPLEMENTAL DISCUSSION

#### Sex chromosome identification and delimitation of the non-recombining region

We built our initial W-mer list (female-specific *k*-mers) using Illumina whole-genome sequencing data from a single F1 mapping population (Table S9). These isolates were phenotyped for flower sex and 16 were female (ZW) and 17 were male (ZZ) [1]. We used Jellyfish v2.3.0 [2] to identify all *k*-mers in an isolate and used the bash *comm* command to generate a list of *k*-mers shared in all female isolates and not found in any male isolate (W-mers). We mapped the W-mers using BWA-MEM v0.7.17 [3], (with parameters '-k 21', '-T 21', '-a', '-c 10') and found the W-mers densely clustered to Chr09 at ~44.32-47.26 Mb of haplotype 1 (HAP1), with some lower-density, discontiguous peaks flanking this region. This supports the identity of this chromosome as the W and this region as the non-recombining, female-specific SDR.

Because this approach relies on identifying *k*-mers found in one sex, but not the other, we expect that increased genetic distance of isolates may reduce the W-mer signal due to *k*-mers from the SDR matching autosomal *k*-mers. To test the effect of different sampling strategies on SDR identification we built three additional W-mer lists. Previous analyses showed that genetic variation in *Amborella* was structured into four geographic regions across New Caledonia [1]. We sampled within populations for Aoupinie (13 females, 17 males) and Dogny (11 females, 11 males), representing two distinct population clusters. We also used *Amborella* genotypes collected from across the geographic range (referred to as the Island-wide sampling; six females, nine males) [1] and compared these to the mapping population. Together, these additional sampling strategies, that span the geographic range of *Amborella,* support a "core" SDR boundary at ~44.32-47.26 of Chr09 in HAP1 (Fig. 2, S4-6).

Some variation around the "core" boundary is observable depending on sampling strategy, as is the density of W-mers mapping to the region (Fig. 2). However, importantly, all four W-mer lists share the same region of Chr09 in HAP1 (Fig. S3-6) with little and sporadic coverage across the other regions of the genome (Fig. S4-6) that could be due to noise from transposable element variation, population structure, and potential sex determination region (SDR) boundary expansion and contraction variation. It is unclear the extent to which each of these sampling strategies would work in other species without an *a priori* understanding of the heterozygosity, population structure, repeat content, and ploidy, as each of these may affect the identification of sex-specific *k*-mers. Therefore, selection of the isolates used should be carefully considered when generating sex-specific *k*-mer lists in a previously unexamined species. Moreover, we did not take into consideration the effect of depth of coverage of the Illumina data, which could additionally affect density of W-mers identified.

The W-mer approach enables the identification of the SDR using fewer individuals than other standard approaches for sex chromosome identification, such as a linkage-map. Here we found that using as few as three individuals of each sex permitted identification of the SDR in *Amborella*, although noise along the autosomes decreases as more individuals are used (Fig. S4-6). In total, these findings illustrate the utility of sex-specific *k*-mers in delimiting the SDR boundary with the PAR, even in a homomorphic sex chromosome system with a relatively small SDR (~3 Mb) and limited divergence (mean Ks =0.0298, SD=0.032; Fig. 3).

While *k*-mers have been previously used to discover sex-linked sequences [4,5], to our knowledge this approach has not been readily used for PAR/SDR delimitation. To validate our findings when using W-mers, we additionally used standard approaches of sex-specific read coverage and nucleotide diversity between the sexes to identify the sex chromosomes. Coverage is relatively consistent across the Z and W for both sexes, though a few regions are clearly W-specific in HAP1 (Fig. S9). Read coverage was calculated by mapping the reads to HAP1 and 2 separately using BWA v0.7.17 [3]. Coverage was calculated on the resulting bams in 10,000bp windows using BEDTools v2.28.0 [6]. Because the Z and W reads map well to both haplotypes in the SDR, we calculated the average nucleotide differences between the sexes (dXY; described in the Methods). We identified the same region of Chr09 in HAP1 as when using the W-mers. But this approach identifies the homologous region of the Z (HZR) chromosome in HAP2 (Fig. S9) equally as well. Additional analyses would be necessary to test for phasing of the Z/W pair (e.g., fixed female-specific SNPs) and would likely require the combination of sex-specific read coverage and population genomic analyses to identify highly-diverged or W-specific regions (same logic applies to the Y chromosome in an XY system). This highlights an additional benefit of using sex-specific *k*-mers for sex chromosome identification: readily testing for phasing of the sex chromosome pair. We expect this approach to be similarly useful in other scenarios that involve blocks of suppressed recombination, such as super genes.

#### SUPPLEMENTAL METHODS

##### Identifying karyotypic sex of isolates

An additional use of sex-specific *k*-mers is rapid identification of the karyotypic sex of samples. For the isolates that were collected without flowers present we identified whether they contained evidence for a W chromosome. To accomplish this, we used the W-mers in all isolates of mapping population to identify W-linked reads. We used the *bbduk* function of BBmap v38.86 (Bushnell, sourceforge.net/projects/bbmap/) to identify reads that contained perfect matches to the list of W-mers. We mapped these reads to the W-containing HAP1, using BWA v0.7.17 [3] and used Samtools v.1.10 [7] to calculate coverage of the non-recombining region, using the region at 44.32-47.26 Mb. We plotted these coverage values using ggplot2 [8] and found a clear binary pattern. Consistent with the expected 1:1 sex-ratio found in *Amborella* [9], we found about half of the individuals show similar coverage in the SDR as the female genome line. We consider these as females and the individuals with nearly no coverage in the SDR males.

##### Construction of the scaffold assembly

A total of 3,605,703 PacBio reads (58.81x per haplotype) were assembled using HiFiAsm+HIC assembler [10], and formed the starting point of the version 2.0 release. The 158,007,088 Illumina fragment 2x250 reads (49.62x sequence coverage) was used for fixing homozygous snp/indel errors in the consensus. Chromosomes were scaffolded using the 374,400,434 2x80 (42.31x) Hi-C reads.

##### Screening and final assembly releases

Scaffolds that were not anchored in a chromosome were classified into bins depending on sequence content. Contamination was identified using blastn against the NCBI non-redundant nucleotide collection (NR/NT) and blastx using a set of known microbial proteins. Additional scaffolds were classified in the version 2 HAP1 release as repetitive (>95% masked with 24mers that occur more than 4 times in the chromosomes) (70 scaffolds, 8.4 Mb), redundant (unanchored sequence with >95% identity and >95% coverage within a chromosome) (5 scaffolds, 190.7 Kb), chloroplast (444 scaffolds, 29.6 Mb), prokaryote (180 scaffolds, 11.0 Mb), fungal (5 scaffolds, 369.5 Kb), and mitochondria (767 scaffolds, 45.9 Mb). Resulting final statistics for the HAP1 version 2 release are shown in Table S12.

Additional scaffolds were classified in the version 2 HAP2 release as repetitive (>95% masked with 24mers that occur more than 4 times in the chromosomes) (51 scaffolds, 6.6 Mb), redundant (unanchored sequence with >95% identity and >95% coverage within a chromosome) (13 scaffolds, 1.6 Mb), chloroplast (329 scaffolds, 24.5 Mb), prokaryote (110 scaffolds, 7.0 Mb), fungal (1 scaffold, 124.4 Kb), and mitochondria (500 scaffolds, 30.7 Mb). Resulting final statistics for the HAP2 version 2 release are shown in Table S12.

#### SUPPLEMENTAL FIGURES

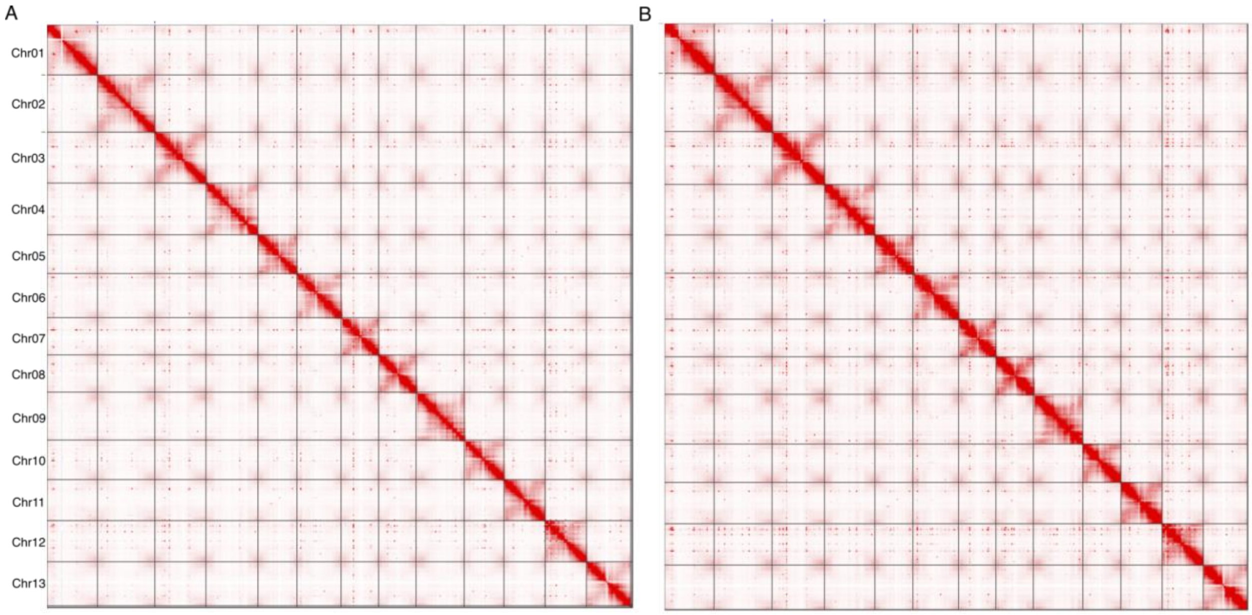


**Fig. S1. Hi-C contact maps for the phased *Amborella* haplotypes.** The Hi-C contact maps show the expected 13 chromosomes for haplotype 1 (A) and haplotype 2 (B).

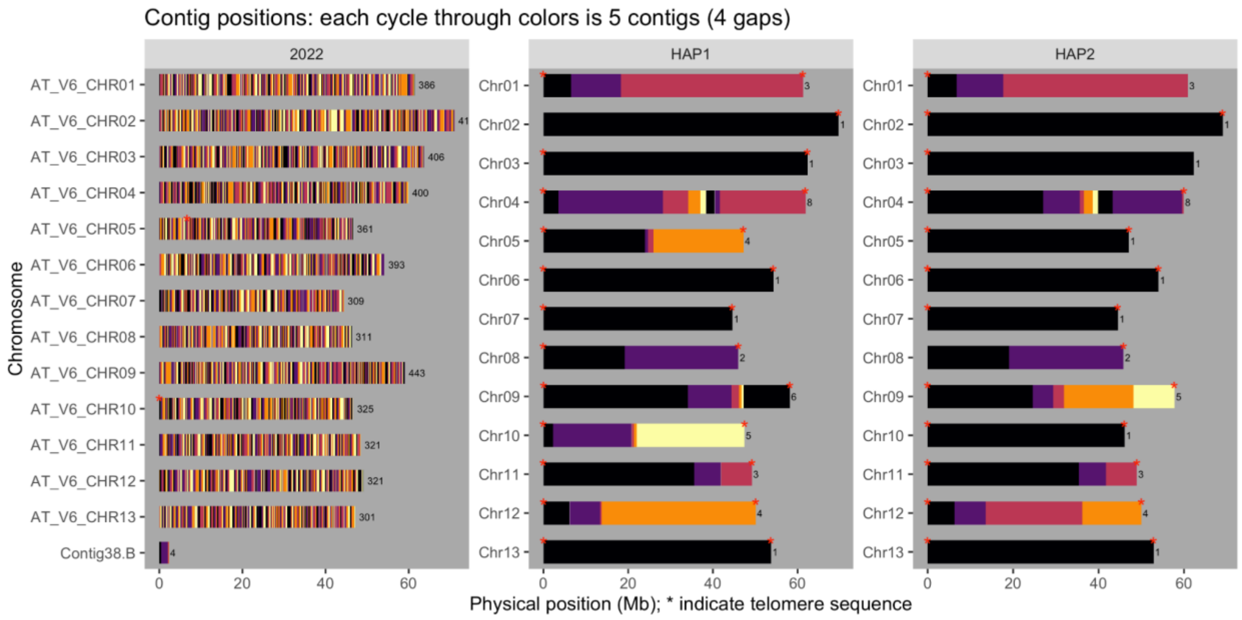


**Fig. S2. Contig map and telomeric sequence location for the *Amborella* genome assemblies.** Each cycle through colors represents 5 contigs (4 gaps). Red asterisks indicate telomere sequences. The v1.0 assembly was not included due being in scaffold-scale rather than pseudomolecules. The contig maps were generated using GENESPACE v.1.3.1 [11].

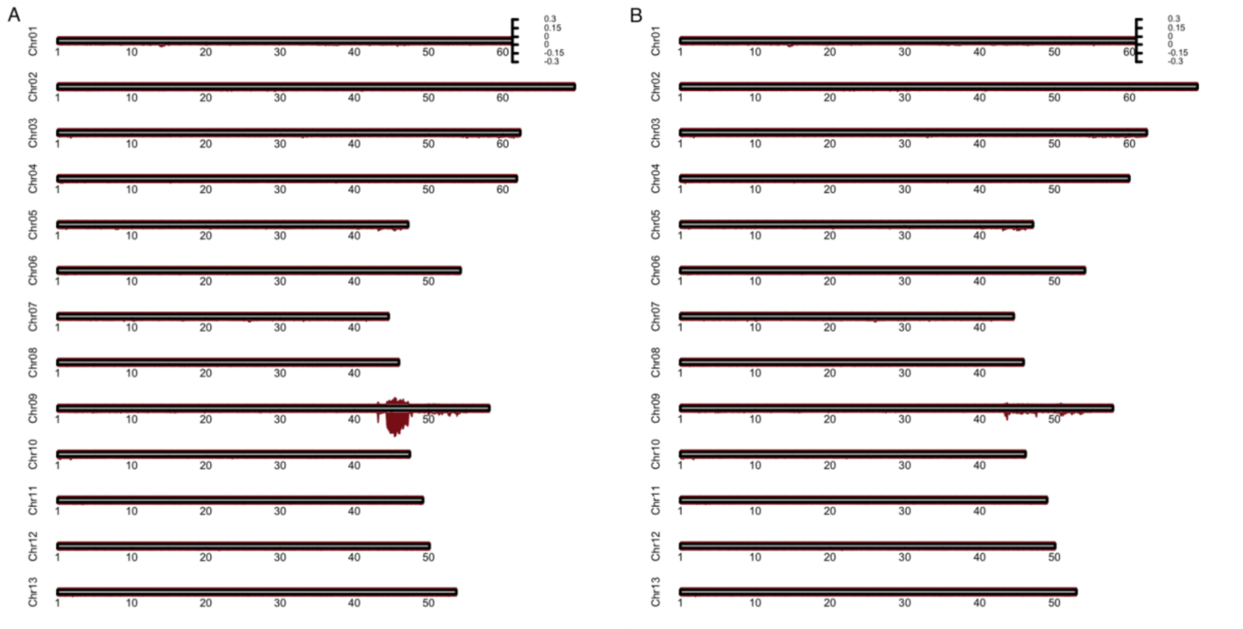


**Fig. S3. Identification of the W chromosome using the mapping population samples.** W-mer coverage across all chromosomes in haplotype 1 (A) and 2 (B). The positive axis shows W-mer coverage when using all isolates from the mapping population (16 females, 17 males). The negative axis shows when using only three of sex.

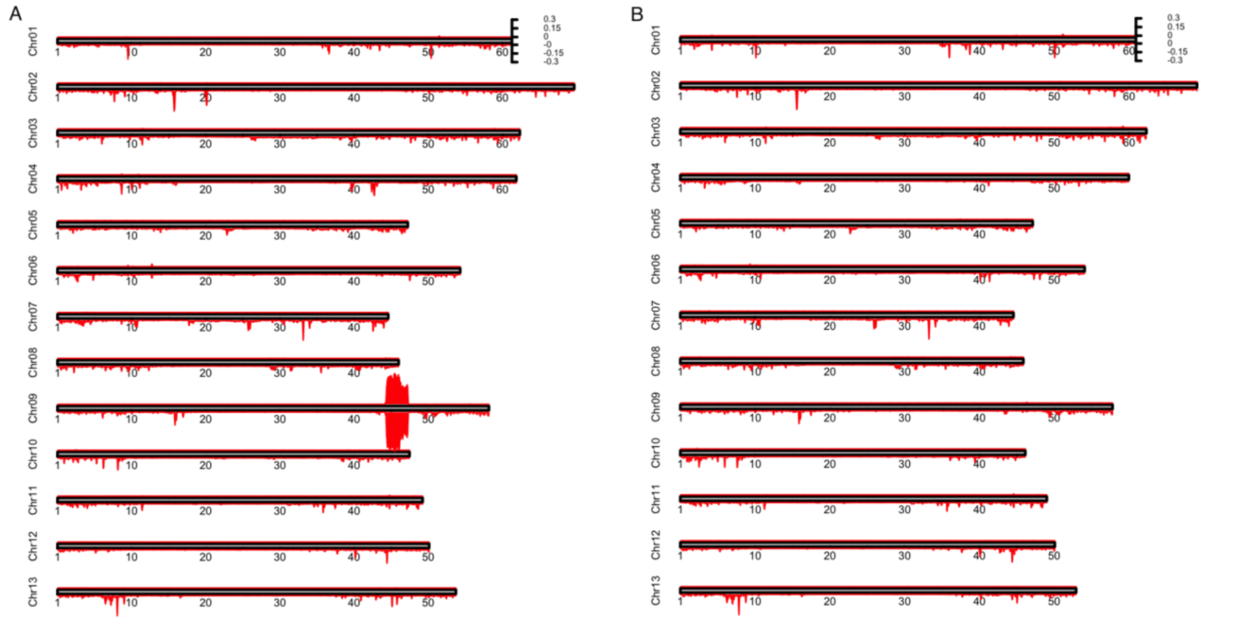


**Fig. S4. Identification of the W chromosome using the Island-wide samples.** W-mer coverage across all chromosomes in haplotype 1 (A) and 2 (B). The positive axis shows W-mer coverage when using all isolates from the island-wide samples (6 females, 9 males). The negative axis shows when using three of sex.

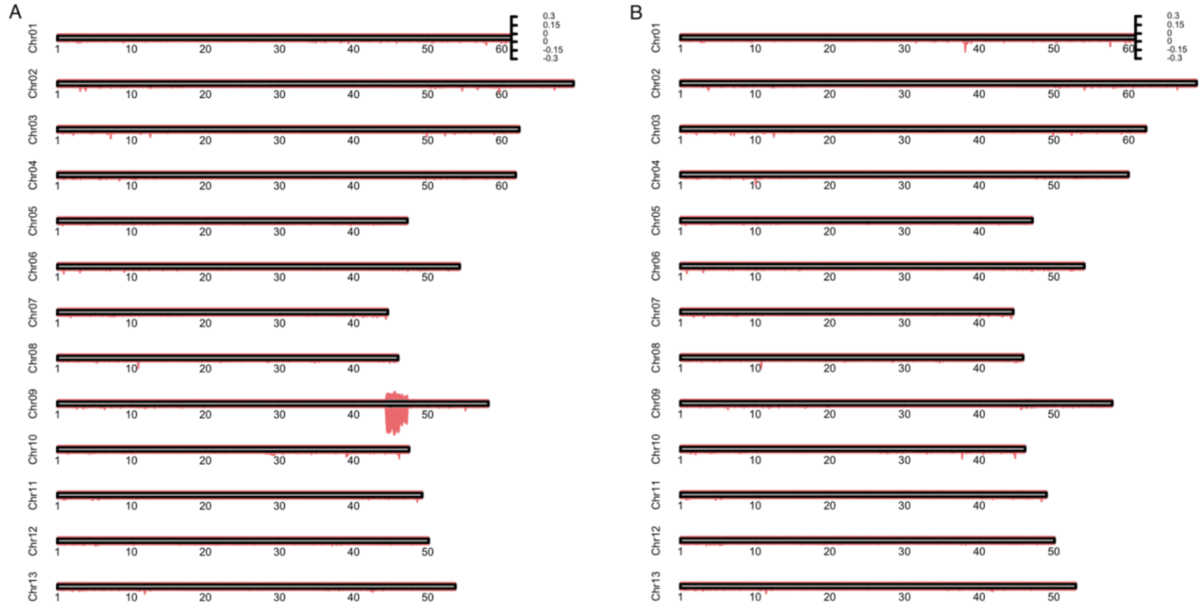


**Fig. S5. Identification of the W chromosome using the Aoupinie samples.** W-mer coverage across all chromosomes in haplotype 1 (A) and 2 (B). The positive axis shows W-mer coverage when using all isolates from the Aoupinie population (13 females, 17 males). The negative axis shows when using three of sex.

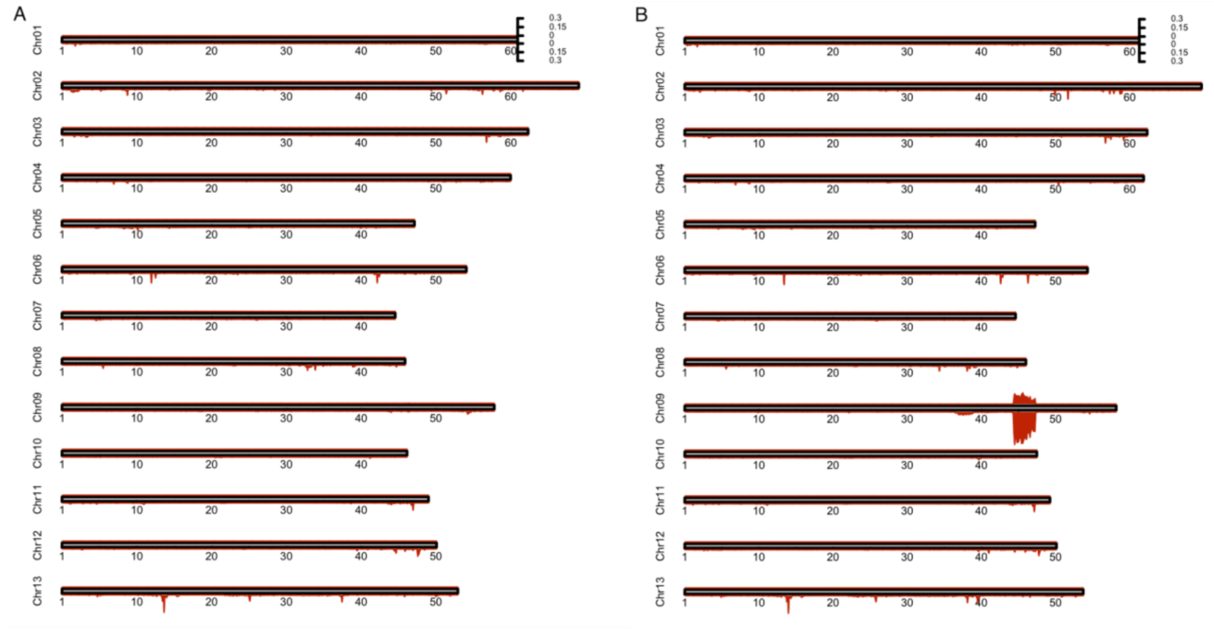


**Fig. S6. Identification of the W chromosome using the Dogny samples.** W-mer coverage across all chromosomes in haplotype 1 (A) and 2 (B). The positive axis shows W-mer coverage when using all isolates from the Dogny population (11 females, 11 males). The negative axis shows when using three of sex.

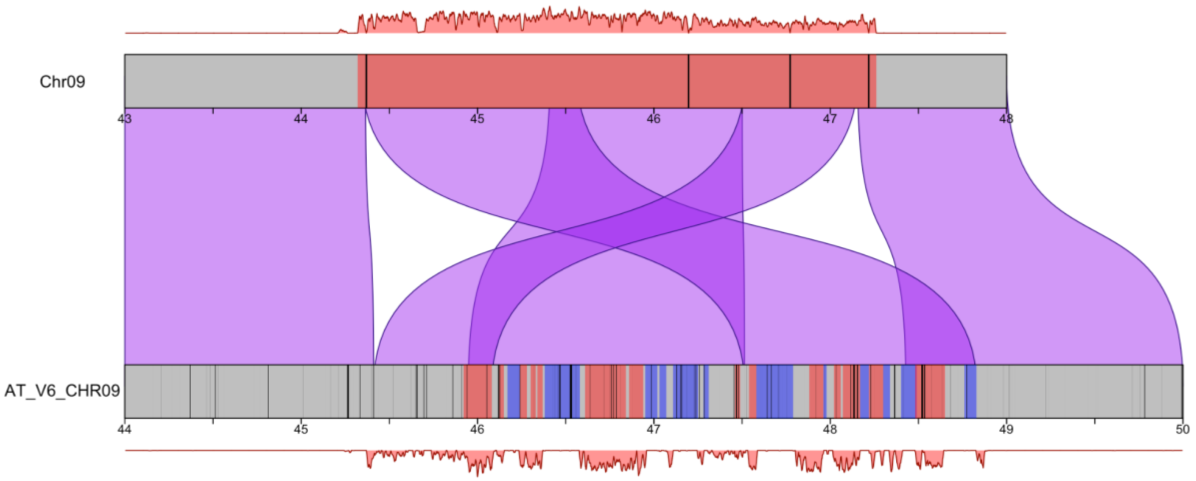


**Fig. S7. Comparison of the phased *Amborella* sex-determining region to a chimeric assembly.** Coverages shown in red are W-mers when using all isolates from the Island-wide sampling. Red boxes in the chromosome ideograms highlight identified W-linked regions and blue Z-linked. For the AT_V6_CHR09 chromosome, the Z vs W-linkage were previously identified in [12] Black lines indicating gaps in the assembly. The purple bars between chromosomes show syntenic blocks identified using GENESPACE [11].

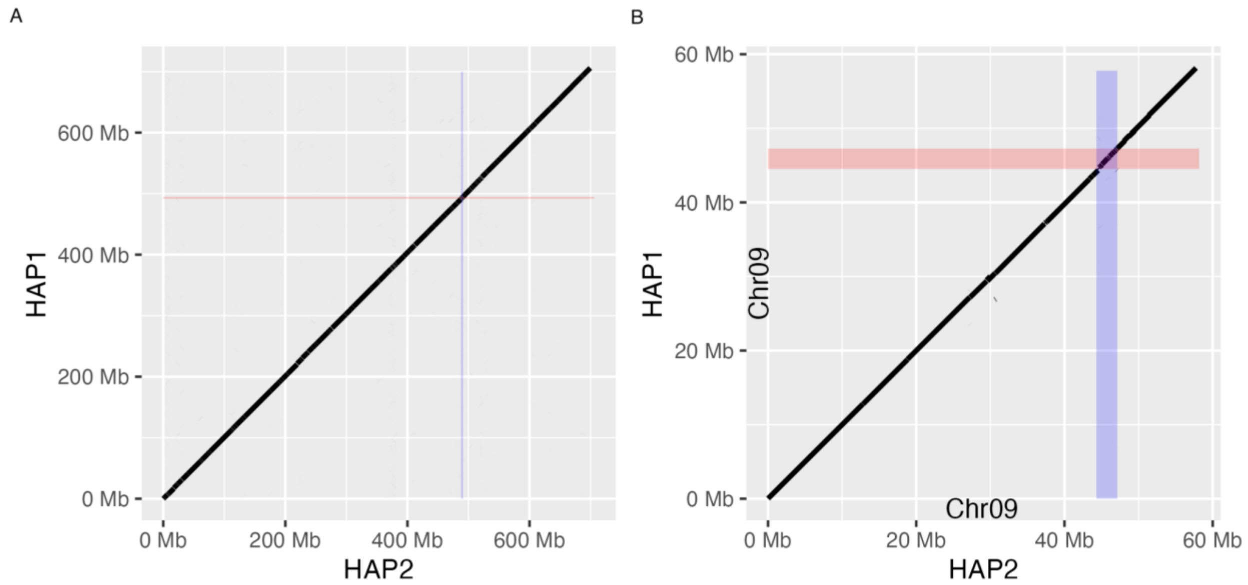


**Fig. S8. Using genome alignment to identify the homologous region of the Z sex chromosome to the W sex-determining region.** A) Dot plot for all chromosomes. B) Dot plot for Chr09 only; the W (HAP1) and Z (HAP2) sex chromosomes. The red highlighted region indicates the sex-determining region of the W located on Chr09 of HAP1 and blue highlights the homologous region of the Z in HAP2.

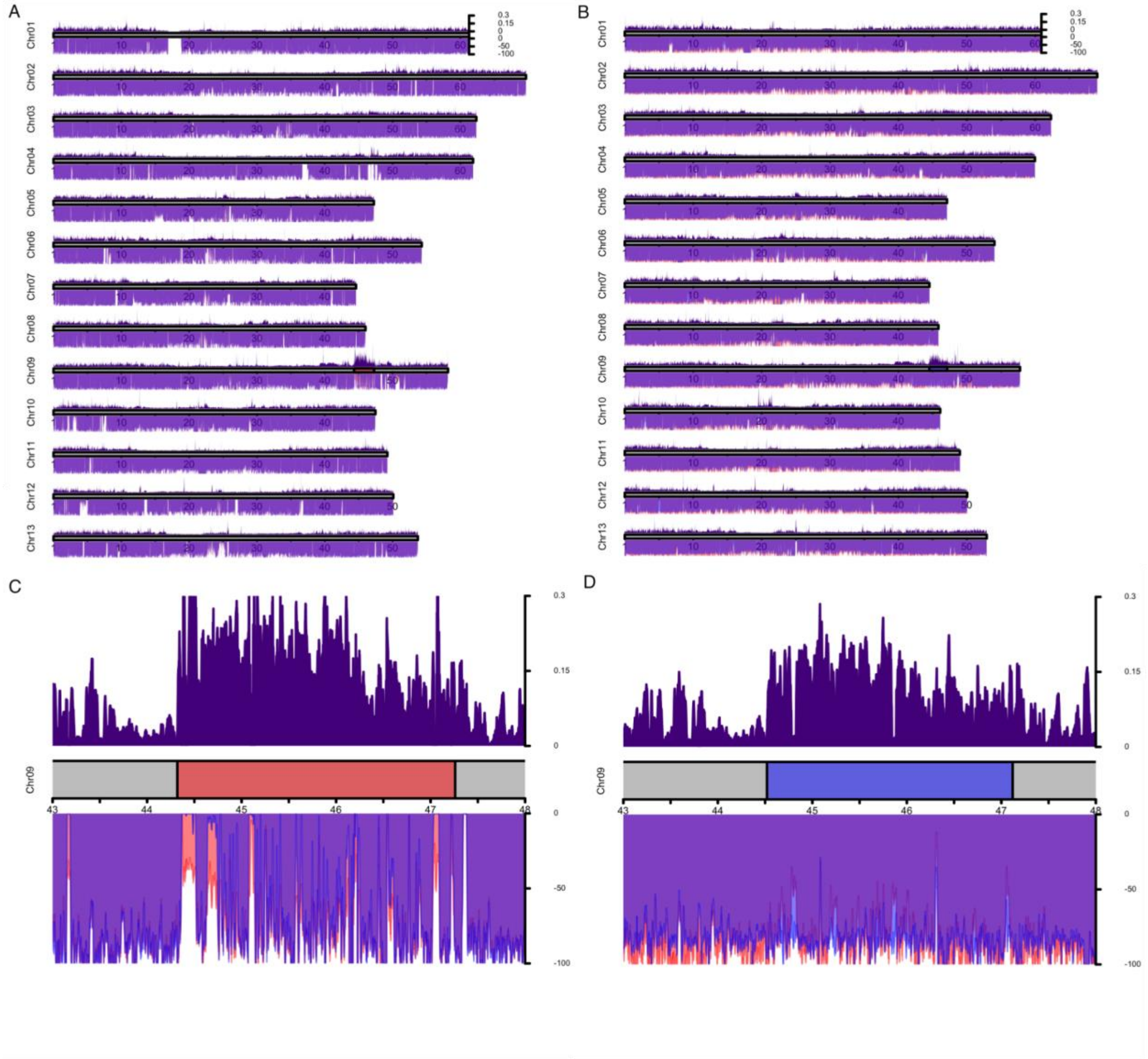


**Fig S9. Sex-biased read coverage and population genomic analyses for comparison of the identification of the sex chromosomes.** The positive axis shows dXY between the sexes (6 females, 9 males) when using the island-wide samples mapped to haplotype 1 (A) and 2 (B). The negative axis shows female (red) and male (blue) Illumina read coverage (capped at 100x) when mapped to haplotype 1 (C) and 2 (D). The red and blue boxes within the ideograms highlight the SDR (A, C) and HZR (B, D) coordinates based on the W-mer analyses. Read coverage and dXY were calculated in 10,000 bp windows and plotted in karyoploteR v1.26.0 [13].

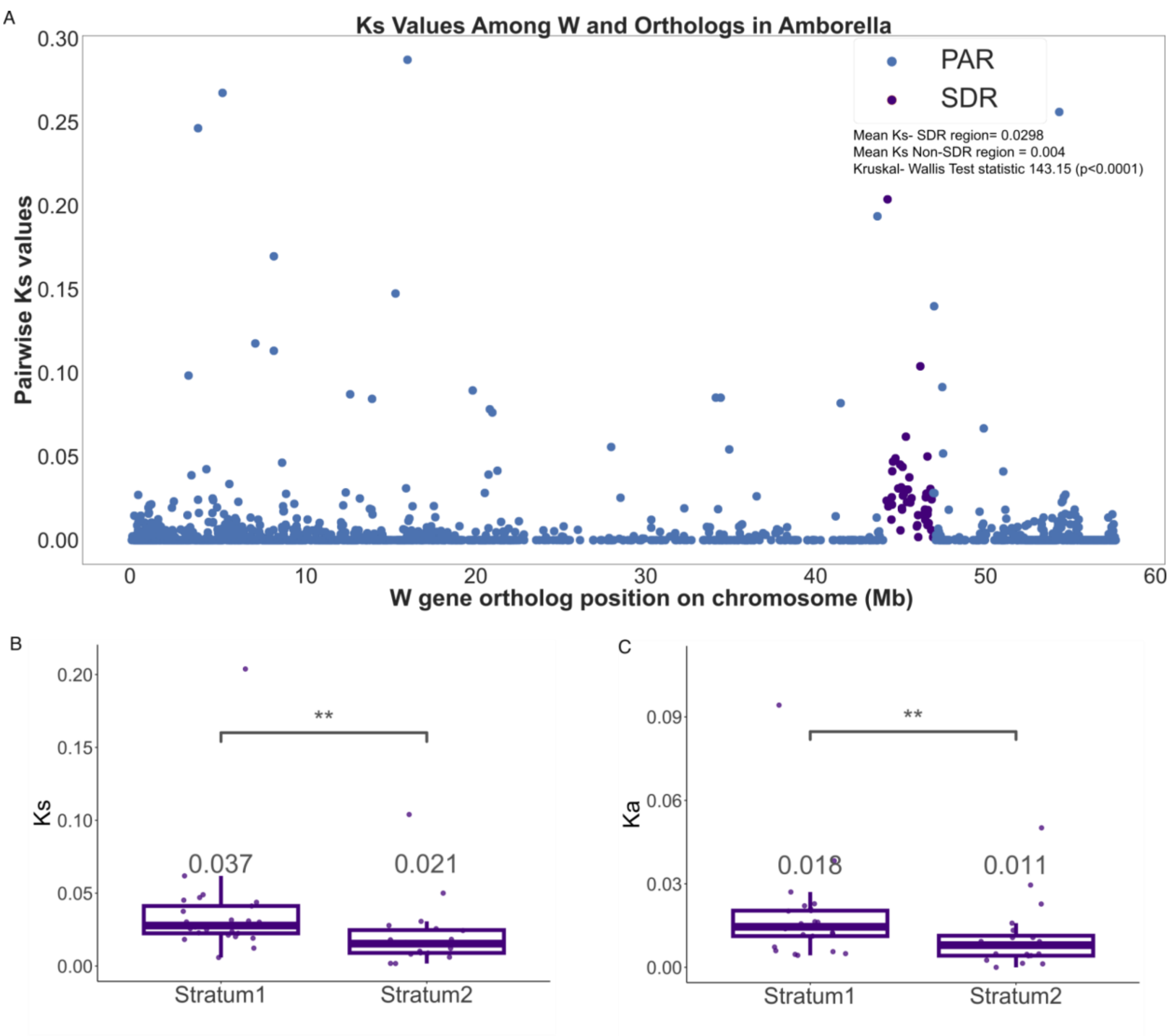


**Fig. S10. Ks analysis on the Z/W chromosomes.** A) Ks across all of Chr09, with gene positions relative to haplotype 1. Purple dots indicate genes located in the sex-determining region (SDR) that have a one-to-one ortholog to the Z, while blue dots are in the pseudoautosomal region (PAR). B) Boxplots of Ks values between the two strata in the SDR and Ka (C). The asterisks indicate the level of significance between strata and the values above the boxes are the average value for that stratum.

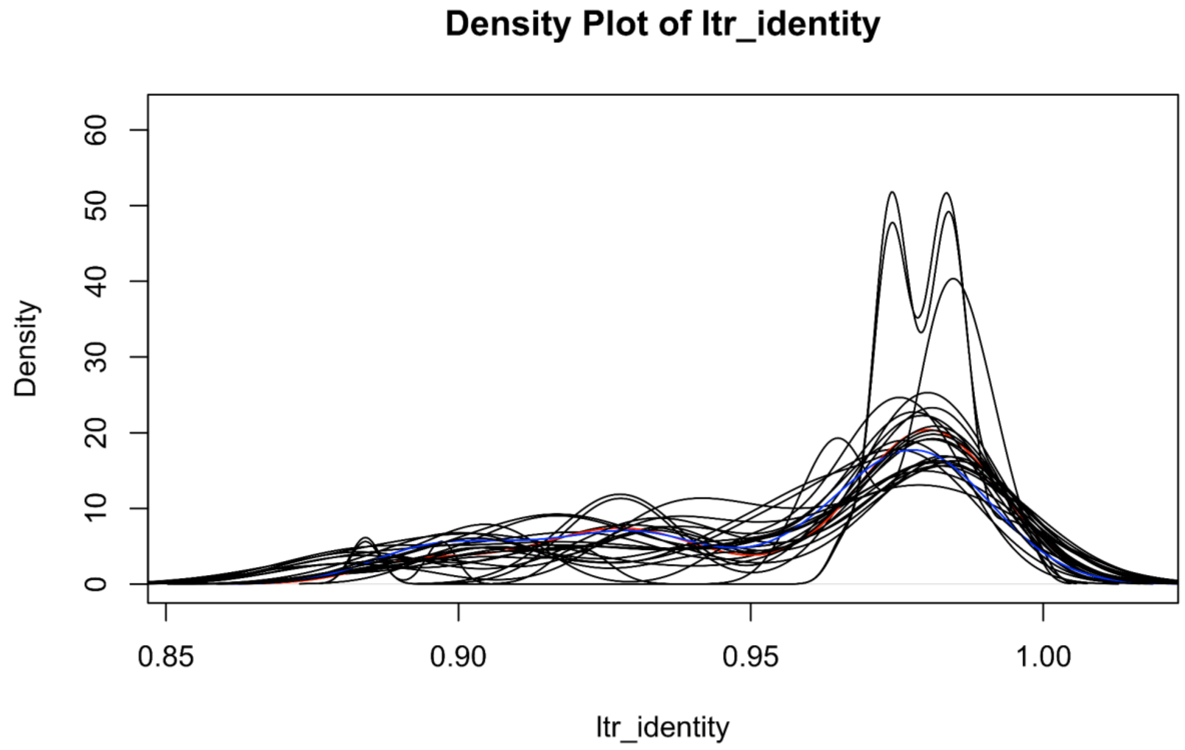


**Fig S11. Density of LTRs by their identity values for each chromosome within *Amborella*.** The density (kernel density estimation) of particular LTR identity values for all 13 homologous pairs. haplotype 1, Chr09 is shown in blue, whereas haplotype 2, Chr09 is shown in red. Their distribution does not differ from the autosomal trend. Homologous pairs tended toward similar kernel densities, such as the bimodal patterns seen between values 0.95-1.00 for Chr02.

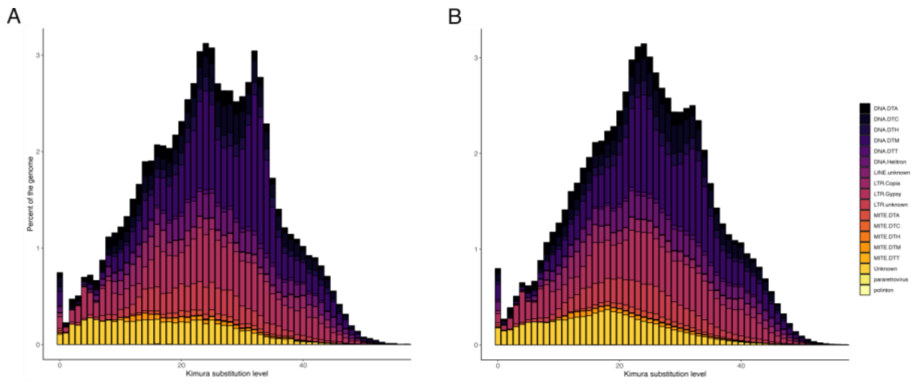


**Fig S12. The repeat landscapes of the *Amborella* sex chromosomes.** A) The W chromosome (Chr09 of haplotype 1). B) The Z chromosome (Chr09 of haplotype 2). Relative time is determined by the Kimura substitution level with lower values closer to 0 representing more recent events and higher values approaching 40 representing older events.

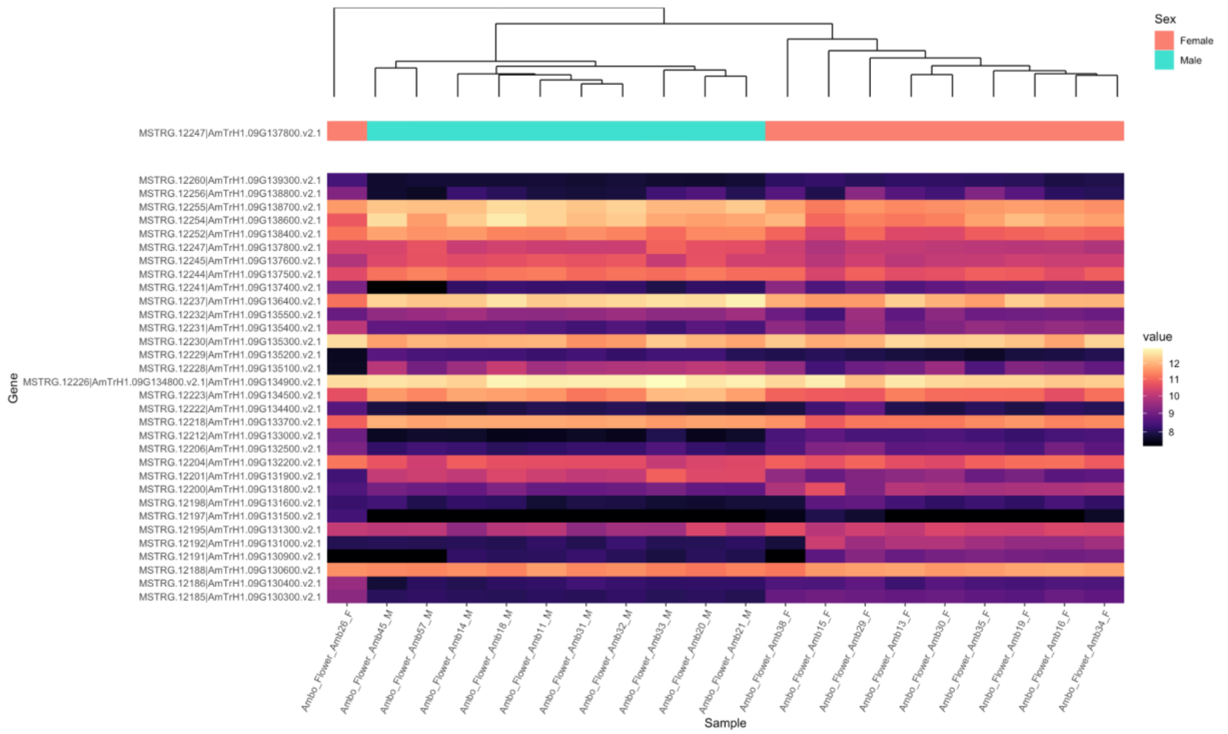


**Fig. S13. Heatmap of significantly differentially expressed genes.** Genes shown were significantly different between females and males of *Amborella* flowers at stage 5/6 at an adjusted p-value less than 0.05 and found in the sex-determining region.

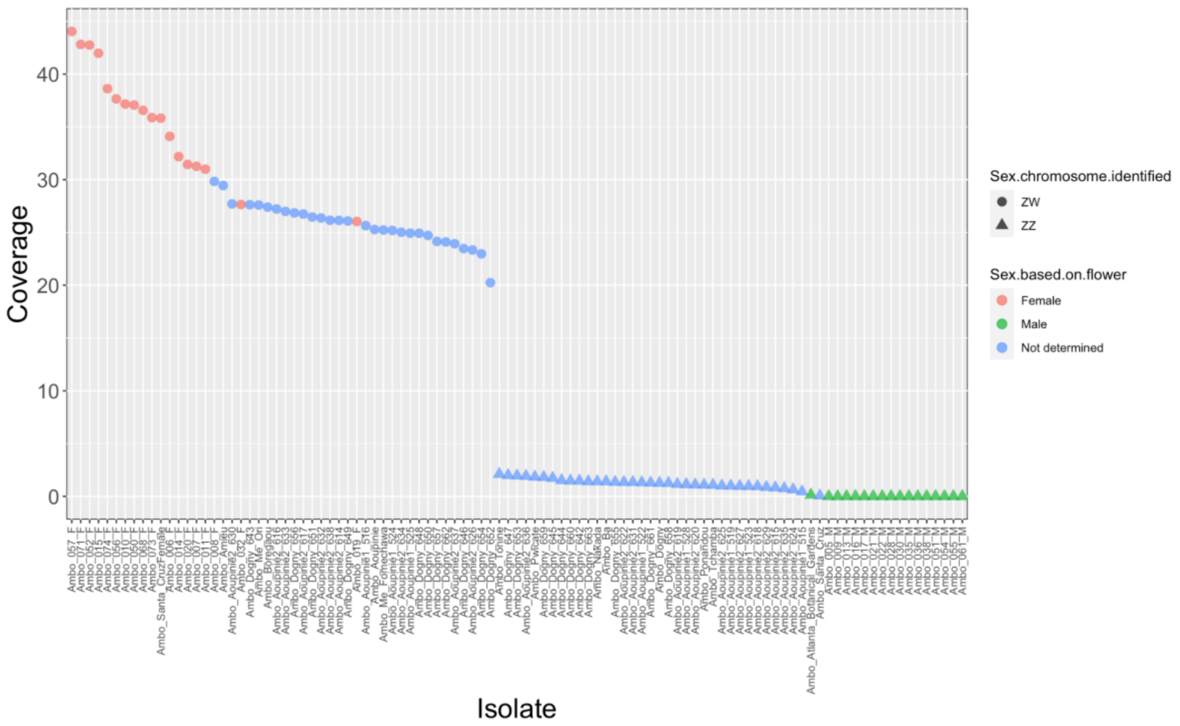


**Fig S14. Identifying karyotypic sex of isolates.** Isolates identified as female on left (circles), males (triangles) on right. The asterisks indicate two isolates of known sex, who were the parental lines of the mapping population isolates that were used to generate the W-mers. These lines were not phenotyped for sex, so we leveraged the W-mer list from the mapping population samples to identify genotypic sex. We identified the reads with exact matches to the W-mers and mapped these back to the HAP1 (W) reference and calculated coverage in the SDR. We found a clear presence-absence pattern, where only genotypic females harbored W-mers.
